# Understanding Molecular Links of Vascular Cognitive Impairment: Selective Interaction between Mutant APP, TP53, and MAPKs

**DOI:** 10.1101/2023.12.08.570506

**Authors:** Melisa Ece Zeylan, Simge Senyuz, Ozlem Keskin, Attila Gursoy

## Abstract

Vascular cognitive impairment (VCI) is an understudied cerebrovascular disease. As it can result in a significant amount of functional and cognitive disabilities, it is vital to reveal proteins related to it. Our study focuses on revealing proteins related to this complex disease by deciphering the crosstalk between cardiovascular and cognitive diseases. We build protein-protein interaction networks related to cardiovascular and cognitive diseases. After merging these networks, we analyze the network to extract the hub proteins and their interactors. We found the clusters on this network and built the structural protein-protein interaction network of the most connected cluster on the network. We analyzed the interactions of this network with molecular modeling via PRISM. PRISM predicted several interactions that can be novel in the context of VCI-related interactions. Two mutant forms of APP (V715M and L723P), previously not connected to VCI, were discovered to interact with other proteins. Our findings demonstrate that two mutant forms of APP interact differently with TP53 and MAPK’s. Furthermore, TP53, AKT1, PARP1, and FGFR1 interact with MAPKs through their mutant conformations. We hypothesize that these interactions might be crucial for VCI. We suggest that these interactions and proteins can act as early VCI markers or as possible therapeutic targets.

## Introduction

Vascular cognitive impairment (VCI) is a severe health issue with cognitive implications and encompasses a broad spectrum of cognitive disorders [1]. VCI causes a significant burden to people and healthcare systems and is the second leading cause of dementia [2]. Therefore, finding early biomarkers and pathways associated with VCI is crucial. VCI is a heterogeneous disease related to vascular and cognitive disorders. Such complex diseases can be analyzed and studied with a systems biology approach. One can study VCI by studying the crosstalk between cardiovascular and cognitive diseases (CVD and CD) by building networks specific to these diseases.

*Network medicine* is a field that focuses on studying diseases with protein-protein interaction (PPI) networks. Most PPI networks are built as static such that each node represents one protein, and their link demonstrates an interaction between the proteins. Nevertheless, proteins are dynamic, and they can adopt several conformations structurally. Therefore, enriching the nodes of the PPI networks with their alternative conformations can unravel the different interactions they can make in the network. The resulting structural PPI network will also contain information about different protein conformations. PRISM [3] is a software for predicting novel interactions using the structural PPI network. PRISM forecasts probable protein interaction using molecular modeling and estimations of binding affinities. Therefore, one can extract more information from a structural PPI network.

Although VCI is a prevalent disease, there is still a substantial lack of understanding of the molecular mechanisms that play a role in the progression of VCI. Our work focuses on finding proteins that can serve as early VCI markers or as potential therapeutic targets. We studied VCI by analyzing the crosstalk between CVD and CD -including possible oxidative stress (OS) implications-. We studied the crosstalk by constructing structural PPI networks related to CVD, CD, and OS. We used PRISM to predict novel interactions in the structural network. PRISM predicted several interactions that can serve as novel VCI-related links. Studying the mutations on the predicted interactions revealed that two mutated versions of APP (V715M and L723P), which were not previously associated with VCI, interact with various proteins. Our results show that two mutated versions of APP have selective interaction with TP53, MAPK3 (ERK1/2) and MAPK14 (p38α). Furthermore, TP53, AK1, PARP1, and FGFR1 also interact with MAPKs through their mutant conformations. We suggest that these interactions might be crucial for VCI.

## Results

### Finding Hub Proteins and Constructing Structural Protein-Protein Interaction Network

We performed a weighted average approach and computed a score for each protein in the Global Networks (GNs) in order to find the hub proteins (See Methods section 1 and 2). We define GN as the merged networks of CVD and CD related to different tissues. **Supplementary Figure 1** and **Supplementary Figure 2** demonstrate the thresholds used for selecting the proteins as hubs. These thresholds also correspond to a score of 1 for each GN. We obtained 57 proteins from the brain GN and 41 from the artery GN. Their sum resulted in 66 unique hub proteins. After including the interactors of the hubs, we ended up with 918 proteins. We incorporated 630 genes refined from GWAS data associated with CVD, CD, and VCI. 27 proteins were both GN and GWAS-related. Five were hub genes (APOE, ATXN1, NEDD4, PARK7, SIRT1). After merging the hubs and interactors with the GWAS genes, we obtained 1521 proteins. We constructed the PPI network with all of these proteins, in which the most connected network resulted in 1438 nodes and 18242 interactions. Including structural information in a PPI network with 1438 proteins would be computationally costly. Therefore, we investigated the most connected cluster of this PPI Network (See Methods sections 3 and 4). The most connected cluster of this PPI Network contains 62 nodes with 1020 edges. Structural information was embedded in this network (See Methods section 5) by integrating 2612 PDBs into the nodes. We used the structural PPI network containing 24085 interactions for molecular modeling with PRISM.

### Protein Modeling with PRISM

PRISM had predictions for 574 interactions with 59 proteins. We assessed the validity of the interactions by investigating their STRINGdb combined score. We examined interactions with combined scores between 0.4 and 0.15 because they have medium to low confidence, making them potentially novel interactions. From the predicted interactions, 168 have a combined score greater than 0.4 (not included). One hundred have a combined score between 0.4 and 0.15; 306 were absent in this scoring (**Fig.1**).

**Fig. 1.**
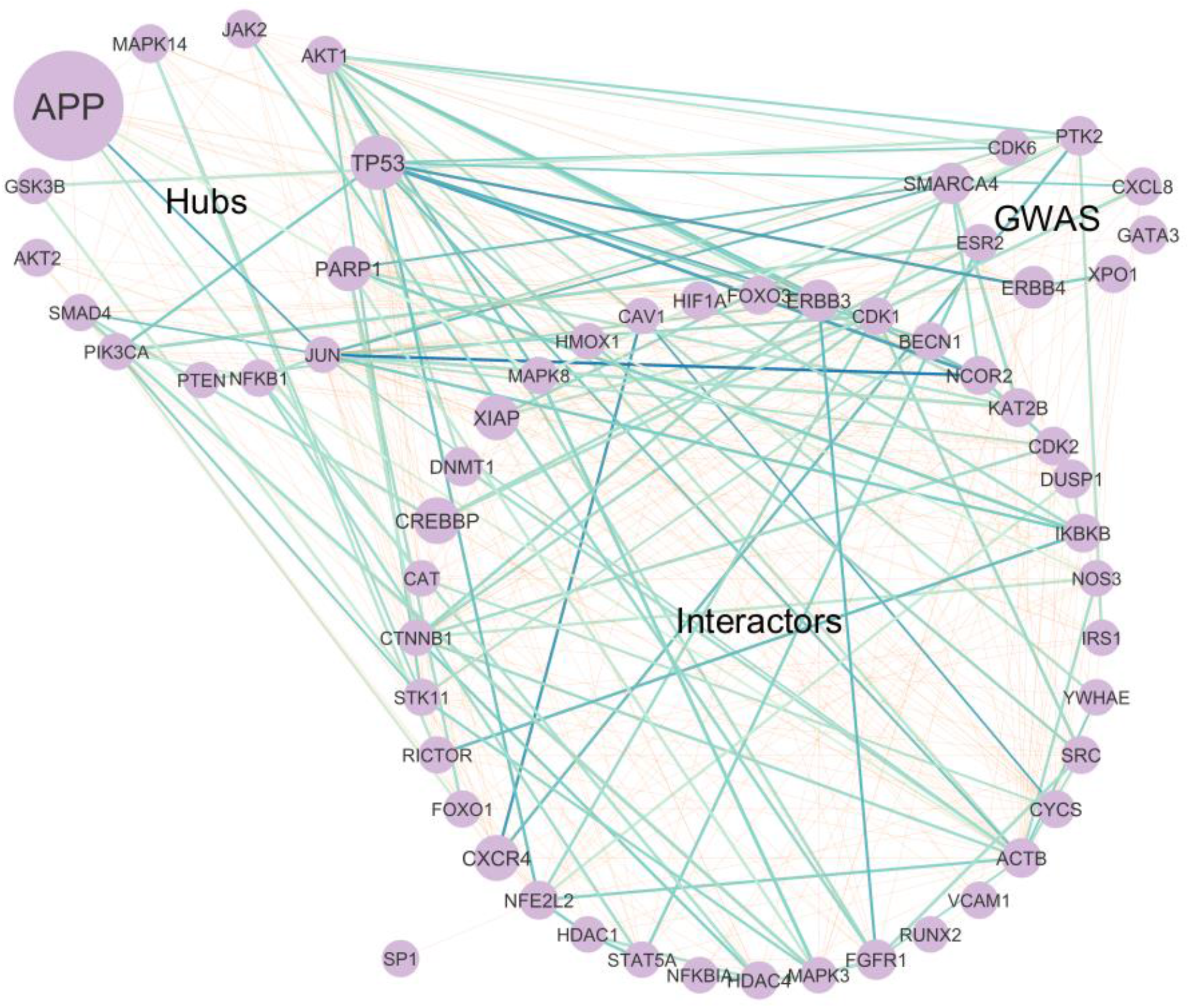
Out of the 574 predicted PRISM interactions, the interactions with a combined score of between 0.4 and 0.15, are demonstrated in shades of blue. The darker shades demonstrate edges with less energy found by PRISM. The rest of the interactions are shown in orange.

We analyzed the locations of the proteins at the cellular level to detect if they were in the same compartment (colocalized). Of the 574 predictions, 413 interactions are colocalized in at least one cellular compartment. The nucleus contained 335 of the interaction predictions; however, some are also in other compartments. The ones not found in the nucleus are mainly in the cytoplasm and can also be in mitochondrion, membrane, and cytoskeleton.

In order to find novel interactions related to VCI, we investigated the PRISM predictions. One hundred sixty-eight predictions (**Fig. 1**) had medium to high confidence. There were 306 that did not have significant experimental or database scores but a combined score of 0.4 from other STRINGdb properties. One hundred interactions had confidence levels ranging from medium to low. Due to their experimental support and PRISM modeling, these 100 interactions may represent novel VCI-related interactions. Out of these 100 interactions of 53 proteins, we surveyed the literature for five interactions with the lowest binding energies (JUN-NCOR2, CXCR4-CAV1, TP53-ERBB4, TP53-NCOR2, and APP-JUN) for their VCI relation. Supplementary Data demonstrates these 100 interactions that are potentially VCI-related. We further investigated if there were mutated structures resulting from the PRISM modellings.

### Mutated protein structures in the predicted interactions

To understand the nature of the interactions found by PRISM, we mapped mutations to the structural PPI network. After mapping the mutations taken from UniProt and ClinVar to PRISM results, several PDB chains are mutated (**Table 1**).

**Table 1.**
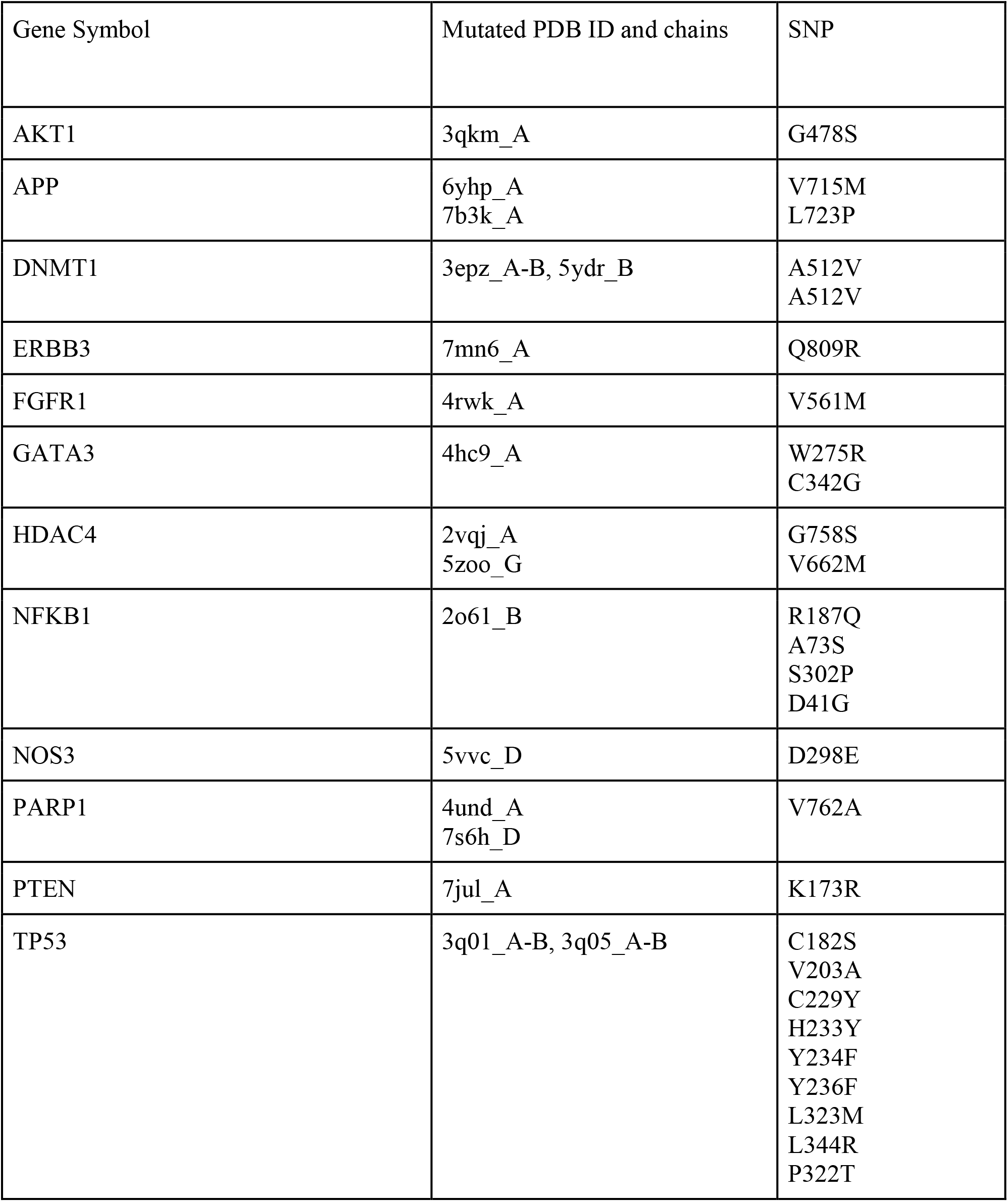
Mutations in the structural network. Number of interactions they make – interactions. For example, for APP: chain A of 6YHP and chain A of 7B3K are mutated. In the table these chains are given with underscore.

From the five putative VCI-related interactions mentioned to be analyzed (JUN-NCOR2, CXCR4-CAV1, TP53-ERBB4, TP53-NCOR2, and APP-JUN), TP53 and APP have interactions predicted with mutated PDB chains (**Fig.2**.). **Fig.2**. demonstrates only the putative VCI-related five interactions unless there is a mutated PDB interaction between the other proteins.

**Fig. 2.**
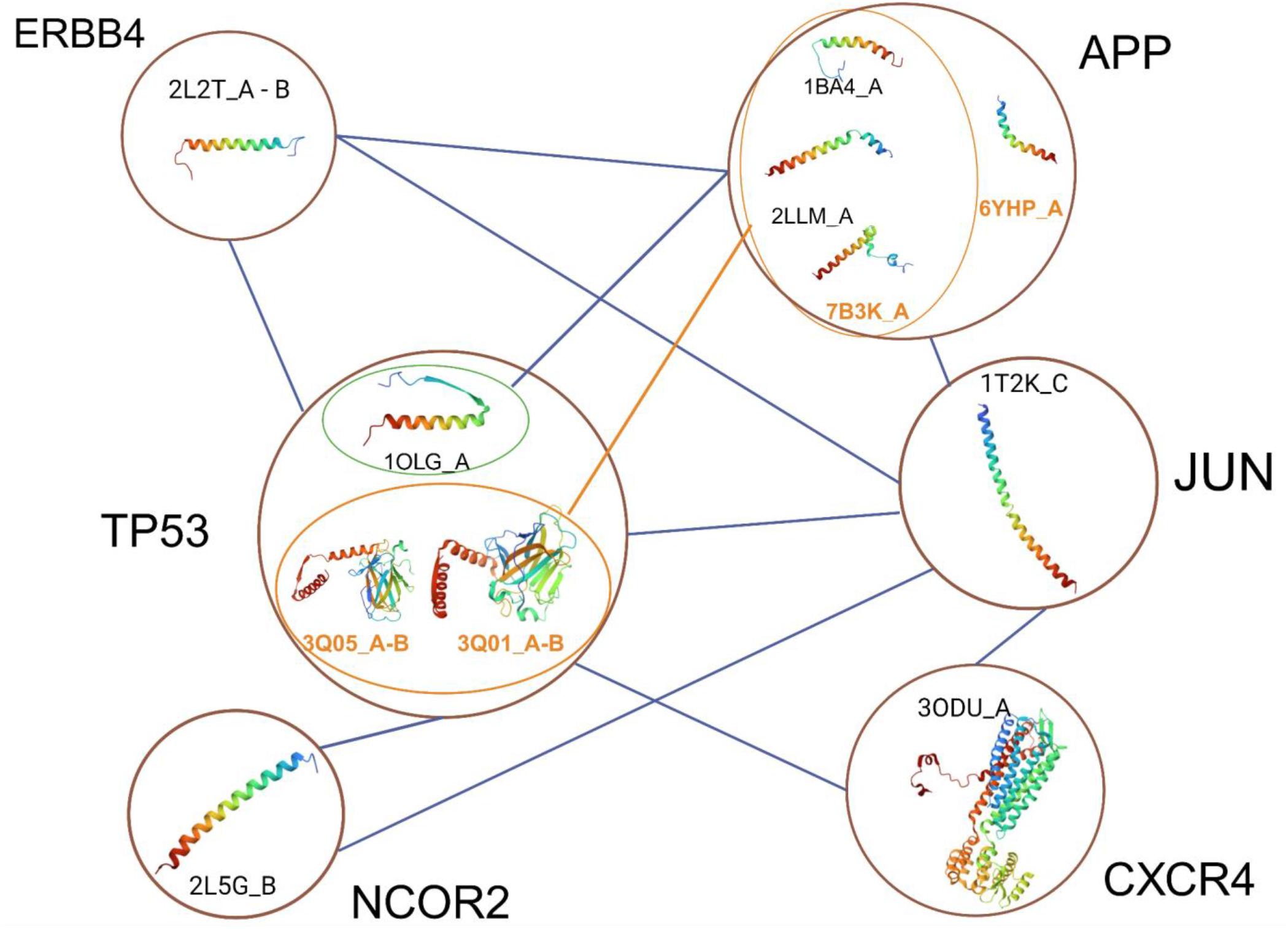
Putative VCI-related interactions and the conformations used for the interactions. Not all the conformations for the given proteins are shown due to visualization purposes. Only the conformations taking place in the putative VCI-related conformations are shown: TP53, NCOR2, ERBB4, CXCR4, JUN, and APP. The labels of the mutated chains are given in orange. The edge is given orange if there is an interaction with a mutated chain.

From these 5 interactions, the given chains of ERBB4, NCOR2, CXCR4, and JUN interact with both the wild type (WT) and mutated chains of TP53 and APP. Furthermore, a WT chain of TP53 interacts with all the APP chains. The mutants of TP53, on the other hand, do not interact with the 6YHP chain, a mutant of APP.

Due to this selective interaction of APP chains, we investigated its other interactions. In total, there are 26 unique interactions of APP (**Fig.3 A**.). In the following interactions, APP uses 6YHP_A, and 7B3K_A as mutated conformations, and many WT conformations which do not have any mutation in UniProt or ClinVar: AKT1, ACTB, BECN1, CAV1, CREBBP, CYCS, ERBB4, HMOX1, JUN, KAT2B, STK11, TP53. On the other hand, MAPK3 does not interact with 7B3K_A, and CDK1, CTNNB1, FOXO3, IRS1, MAPK14, NFKB1, SMAD4, SRC does not interact with 6YHP_A. The following genes interact with only WT APP conformations: CAT, GSK3B, HIF1A, MAPK8, PTEN. It is particularly interesting that all MAPK’s bind to different conformations of APP (**Fig.3 B**.). This demonstrates a particular selective interaction of MAPKs with the mutated conformation of APP.

**Fig. 3.**
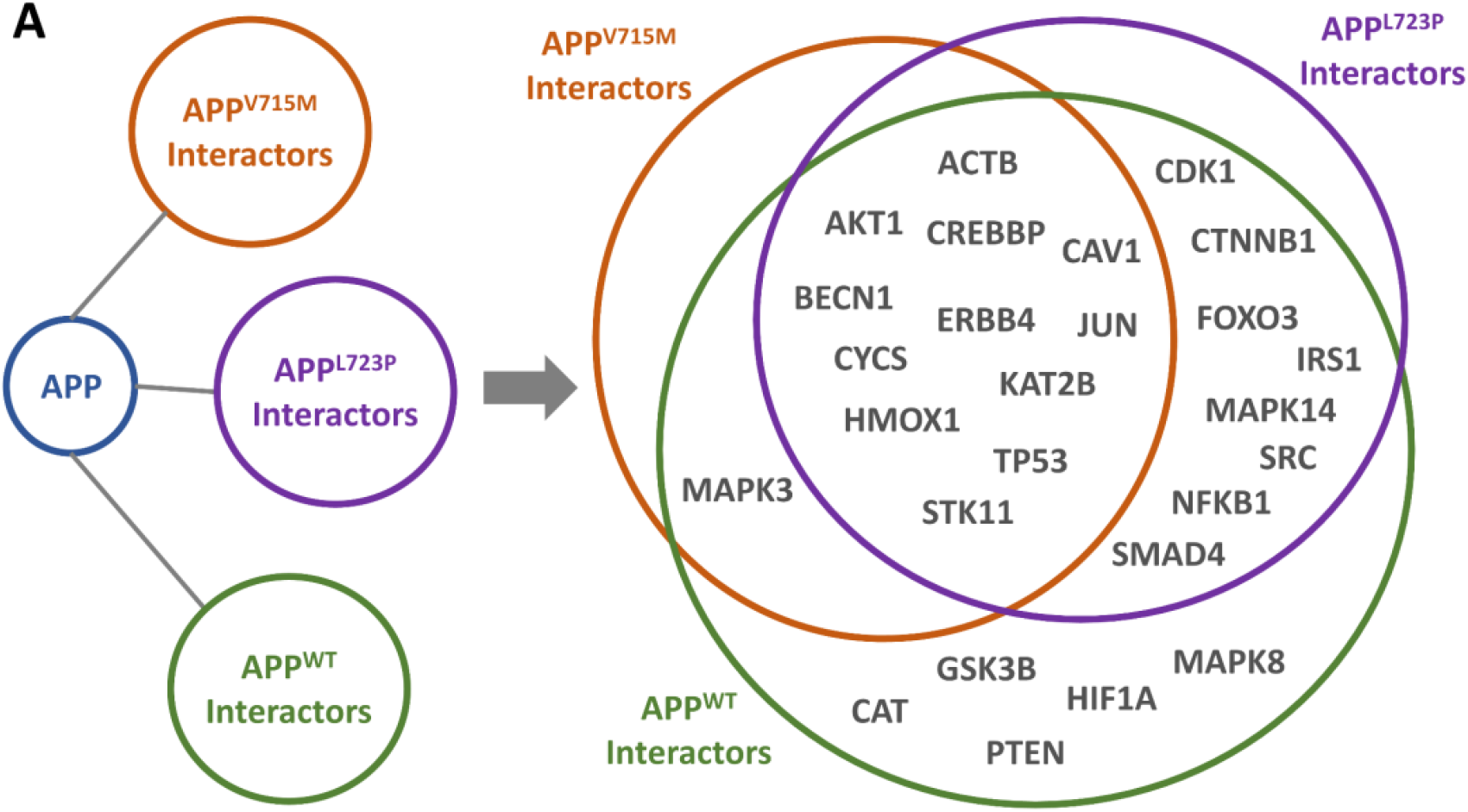
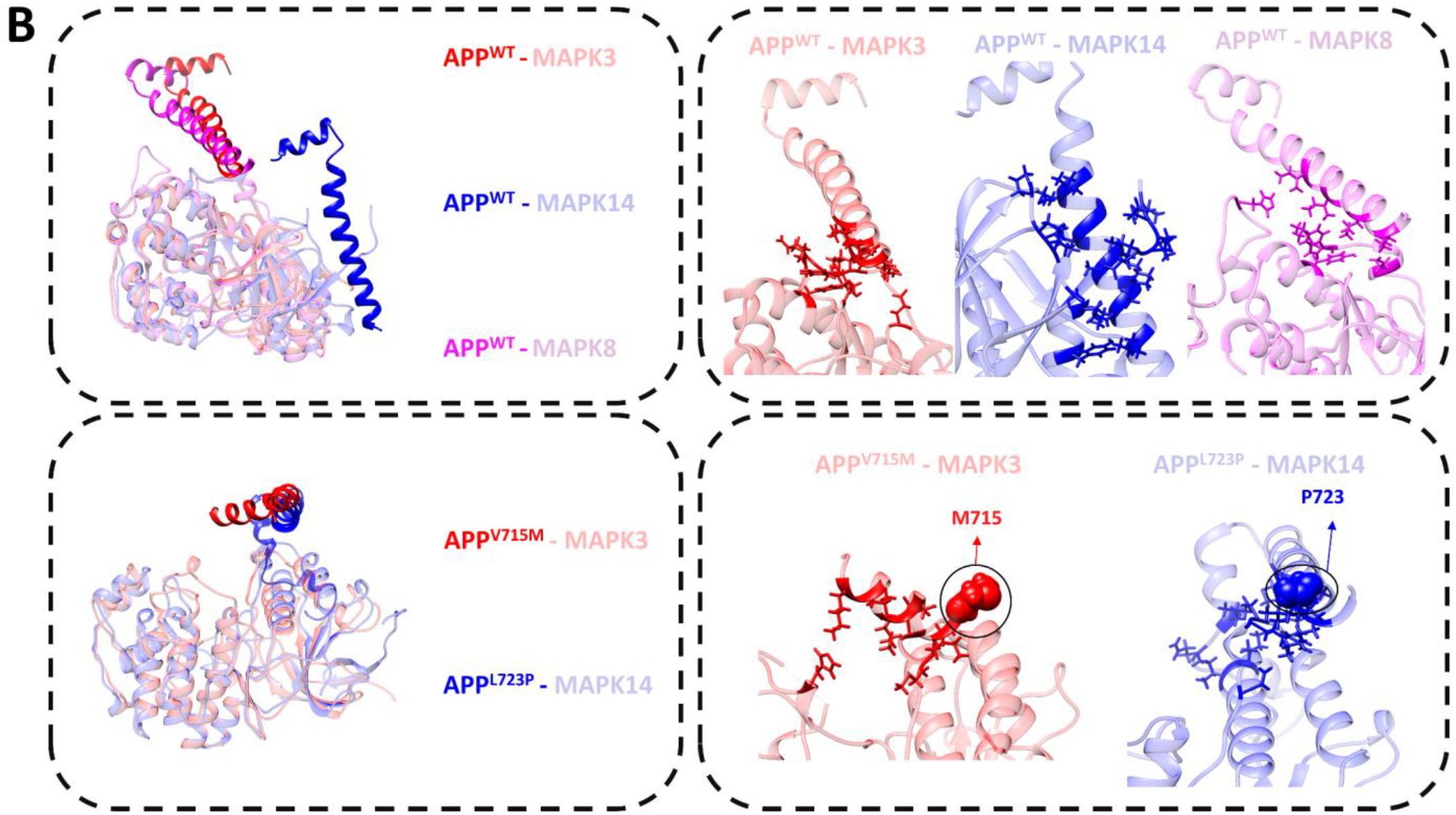
A. Interactors of APP and their preference on 6YHP_A (APP^V715M^), 7B3K_A (APP^L723P^), and APP^WT^ (here, WT stands for any APP conformation which doesn’t have known mutations in UniProt or ClinVar). **B. (left)** Structures of APP^V715M^, APP^L723P^, and APP^WT^ with MAPK14, MAPK3, and MAPK8 are superimposed. APP that binds to MAPK3 (red), MAPK14 (blue), and MAPK8 (magenta) are colored. Corresponding MAPKs are colored as the lighter red, blue, and magenta, respectively. **(right)** Interacting residues are shown as sticks, mutated residues are shown as spheres.

In order to analyze if these MAPKs have a structural preference for other proteins, we included the interactions between the mutated proteins and MAPKs. We discovered that along with APP; AKT1, TP53, and PARP1 also interact differentially with these MAPKs; such that WT or mutated versions of these proteins interact with WT versions of the MAPKs (**Fig.4**.).

**Fig. 4.**
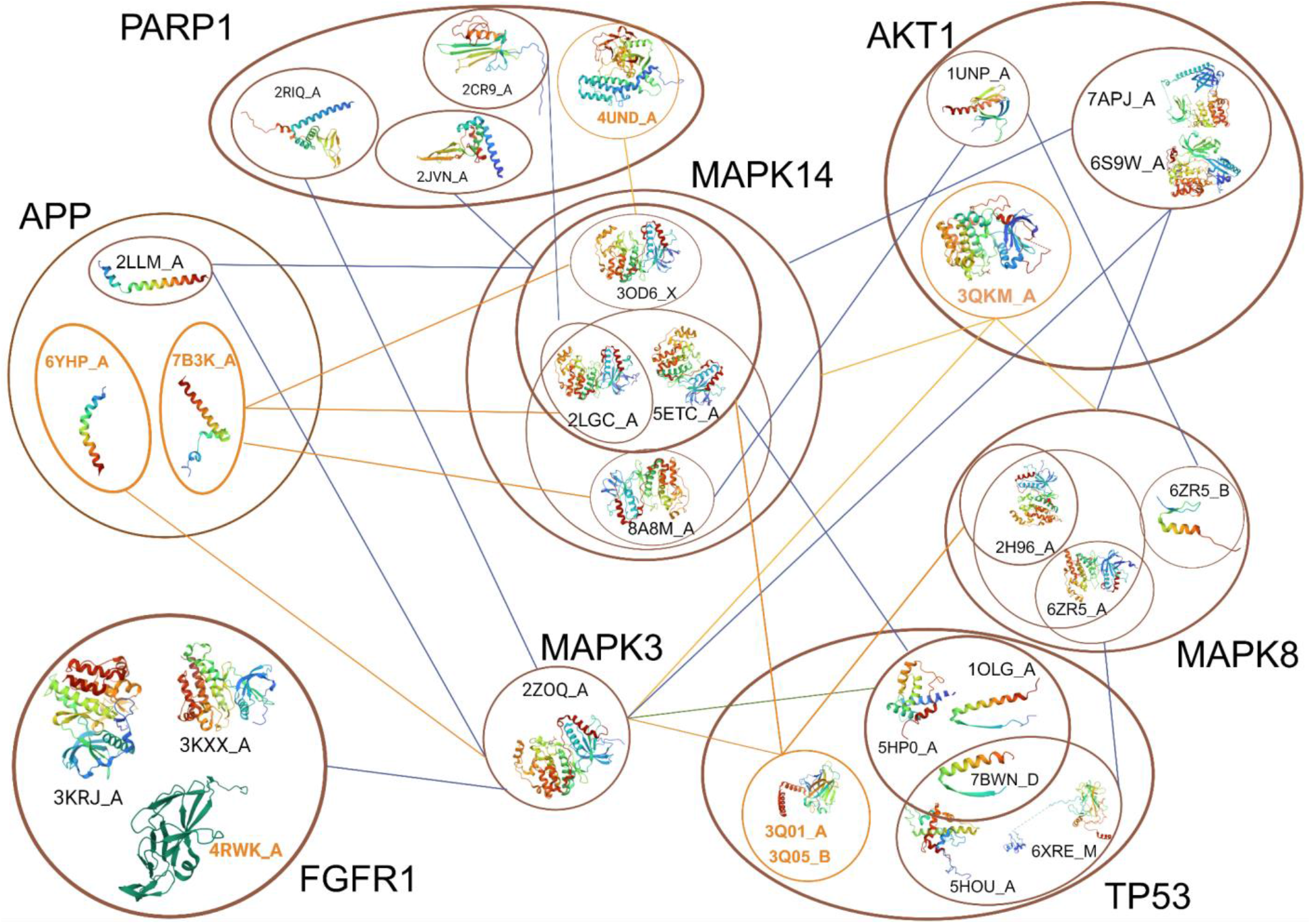
Only interactions concerning the MAPKs were given. Not all the WT interacts are given edges: such that MAPK3 only interacts with one WT of PARP1 (2RIQ_A); we showed this with only one edge representing interaction with the WT. The labels of the mutated chains are given in orange. The edge is given orange if there is an interaction with a mutated chain.

FGFR1 interacts with MAPK3 with all of its conformations. It does not interact with MAPK8 and MAPK14. The mutant chain of PARP1 only interacts with 3OD6_X of MAPK14. And the 2CR9_A conformation has a conformational preference to bind to 2LGC_A. AKT1’s mutant chain can interact with all the MAPKs. The 7APJ and 6S9W structure interacts with all MAPKs. The WT 1UNP chain structure interacts with MAPK14 and MAPK8. This conformation only interacts with the 8A8M structure of MAPK14, and 6ZR5_B structure of MAPK8. TP53’s mutant 3Q01 chain A interacts with all the given MAPKs. It interacts with 2H96 chain A conformation of MAPK8; and, 2LGC_A, 3OD6_X, and 5ETC_A of MAPK14. Additionally, TP53’s mutant chain 3Q05_B only interacts with MAPK14; and further, it only interacts with 2LGC_A, and 5ETC_A conformations of MAPK14. 1OLG_A only interacts with MAPK3 and 2LGC_A of MAPK14. The 5HP0_A chain only interacts with MAPK14 via its 2LGC_A conformation. The 7BWN_D conformation of TP53 interacts with 2LGC_A, 3OD6_X, and 5ETC_A conformations of MAPK14 and 6ZR5_B conformation of MAPK8. The 6XRE_M and 5HOU_A conformation of TP53, also interacts with the 6ZR5_B conformation of MAPK8.

## Discussion

It is essential to understand which proteins can be related to VCI. Our study focuses on finding these proteins by studying VCI with network medicine. With this purpose in mind, we built PPI networks related to CVD and CD and studied their crosstalk. We selected hub proteins and their interactors to refine them for further analysis. Later, we enriched the resulting PPI network with their alternative conformations to incorporate structural information into the PPI network. We later performed molecular modeling on the structural network to discover novel interactions related to VCI. Numerous interactions that may act as PRISM predicted new VCI-related interactions. We investigated if there were mutants within these predicted interactions. Two mutant forms of APP (V715M and L723) that weren’t previously connected to VCI particularly attracted our attention, as TP53 had a conformational preference to one of its mutant conformations. We later discovered that these two mutant forms of APP interact with MAPK3 (ERK1/2) and MAPK14 (p38) with conformational preference. Our results suggest that TP53, AKT1, and PARP1 also have structural preference for MAPKs. These conformational preferences in their interactions might be crucial for VCI.

### Putative VCI-related interactions with protein modeling

In the context of the five putative VCI-related interactions (JUN-NCOR2, CXCR4-CAV1, TP53-ERBB4, TP53-NCOR2, and APP-JUN), we investigated their relations to VCI. In our GNs, the JUN and NCOR2 (SMRT) were present as a hub and an interactor, respectively. In artery tissue, NCOR2 was related to CVD. On the other hand, JUN was in the GNs of both artery and brain tissues connected to CVD. Previous research linked JUN to Alzheimer’s disease and memory loss [4, 5]. Zhou et al. suggested that loss of function in NCOR2 leads to impaired memory [6]. Additionally, previous yeast and mammalian two-hybrid tests [7] demonstrate their interaction (combined score: 0.3). In our results, their binding energy was -34.35. This predicted interaction is potentially localized in the nucleus, where both proteins exist.

The interaction between CXCR4 and CAV1 was present in our GNs as two interactors. CXCR4 was related to a CD in the artery and brain tissue. CAV was present in the GWAS data, and it was also present in the GNs of both brain and artery tissues as CVD-related. Previous research related CXCR4 and CAV1 to CVD and CD [8-11]. Furthermore, with yeast and mammalian two-hybrid tests [12] their interaction was determined (combined score: 0.29), and their binding energy was -29.32. The interaction was colocalized in the me in current literature the interaction between TP53-ERBB4 and APP-JUN is not localized membrane.

TP53 was related to OS in both artery and brain tissues. ERBB4 was related to atrial fibrillation in GWAS data. Previous research detected the interaction of these two proteins with affinity chromatography technology assay [13]. Both were related to cancer in separate studies[14, 15]. Furthermore, the fact that genomic data brought about this interaction highlights the value of including omic data to identify novel disease-related interactions. The interaction of TP53 - NCOR2 [16], and APP and JUN [17] was also previously determined. While the same compartment, TP53-NCOR2 is colocalized in the nucleus.

In conclusion, we discovered three interactions in the same cell compartment —JUN-NCOR2, CXCR4-CAV1, and TP53-NCOR2— out of the five. Even though their likelihood of interacting increased, this does not necessarily indicate an interaction. On the other hand, the other two interactions could still be interacting through intermediary molecules or as a result of mislocalization. To our knowledge, none of these five interactions was discussed in recent literature concerning VCI. We suggest that these five interactions can be novel VCI-related interactions that must be further evaluated.

### Conformational preference of the suggested VCI-related interactions

Understanding the preferred conformations for the putative VCI-related interactions is critical. We examined the structural preferences in these five interactions’ (**Fig. 2**). In these interactions, all the chains interact with each other. Therefore, we can interpret these results as these proteins not having a structural preference when interacting. However, we discovered that TP53 has a conformational preference for a mutant APP structure. Therefore, we first discuss the possible implications of this selective interaction. Regarding the selective conformation interaction, we also discovered that the MAPKs also have a conformational preference for APP (**Fig.3**.). Finally, our results suggested that PARP1, TP53, and AKT1 also have conformational preference for MAPKs.

### Selective Interaction of Mutated APP with TP53

Interestingly, two mutant conformations of TP53, interact with one mutant conformation of APP but not the other. These conformations interact with both WTs. This can point to a structural preference of the mutant TP53s for chain A of 7B3K conformation of APP. The change in the structural preference between TP53 and APP can point to a change in interaction when APP is mutated. This preferential interaction can have implications such that the mutations in APP may have changed the binding specificity or caused a conformational change in APP, causing the exposure of a new binding site. Therefore, a new interaction. From another perspective, the dynamicity and flexibility of APP may change, causing interactions with other conformations of proteins. This type of interaction change can affect APP aggregation in the brain. The protein product of TP53 is p53, which can aggregate to form amyloids [18]. While there is no direct interaction between p53 and amyloid precursor protein, to our knowledge, [19] suggests that the γ-secretase complex, which is involved in generating Amyloid β-peptides, also controls p53-dependent cell death. Further, a study suggested that p53 aggregation or its interaction with tau oligomers may have a role in Alzheimer’s disease [20]. A similar aggregation of p53 or APP might occur due to its selective interaction with mutated APP. We suggest that this selective interaction can be affecting VCI.

### Selective Interaction of Mutated APP with MAPKs

MAPK3 and MAPK14, interact with the same WT structure of APP, and MAPK8 interacts with a different WT structure. In their interactions regarding mutated APP structures, while MAPK3 has a structural preference for the 6YHP chain A, MAPK14 has a structural preference for the 7B3K chain A. Although from the same protein family with MAPK3 and MAPK14, MAPK8 did not interact with these APP chains. Below we discuss the selective interaction of APP with these MAPKs, as APP is one of the few VCI-related proteins (according to GWAS data).

Amyloid precursor protein (APP) is a transmembrane protein [21]. Deposition of the truncated amyloid is a known risk factor for cognitive dysfunction [22]. While the exact mechanism remains unclear, the relationship between APP and vascular cognitive impairment was indirectly shown [23]. Two mutated structures of APP were present in PRISM modelings. V715M [24-26] and L723P [27, 28] are related to familial early-onset Alzheimer’s disease. However, whether these mutations are related to VCI is unknown in current literature. Also, these mutations are not studied in the context of protein-protein interactions of APP to our knowledge.

Our PRISM modelings suggested that APP^V715M^ and APP^L723P^ might have a preference in APP’s interactions. It is interesting to note that, even though they belong to the same family, MAPK3, MAPK8, and MAPK14 interact with different conformations of APP. And these MAPK proteins are colocalized with APP in the cytoplasm (and in the membrane for MAPK3), increasing their chance of interaction. MAPK8 only interacts with WT conformations of APP; MAPK3 and MAPK14 interact with the same WT conformation but different mutated conformations: APP^V715M^ and APP^L723P^ respectively. The selective interaction can be related to conformational changes due to the mutation and therefore structural specificity. The differential interaction can also be related to diverged functions of the two MAPKs when they interact with mutated APPs. Furthermore, the mutations can be causing the exposure or creation of new binding sites that only one of the MAPKs can bind.

MAPK3, MAPK8, and MAPK14, also known as ERK1/2, JNK1-3, and p38, respectively, are mitogen-activated protein kinases (MAPKs), which are essential in controlling a variety of cellular reactions in response to extracellular stimuli [29]. p38 is a kinase activated by stress, and increased levels of reactive oxygen species (ROS) were associated with p38 activation-related neurodegeneration [30]. While growth factors and cytokines activate all ERK1/2, JNK, and p38, only p38 and JNK are activated with stress. Cell proliferation and differentiation are common cellular responses to all of these kinases. Nevertheless, the main difference between JNK and the other two kinases in cellular response is development, stress response, and inflammation [31].

Previous research suggested that amyloid beta activates p38 [32] and it also suggested that RAGE, a receptor for advanced glycation end products, can be a promising candidate for inducing oxidative stress via the p38 pathway [33]. Our results suggest that specific APP mutations can change its interaction with ERK1/2 and p38, exacerbating VCI.

### Selective Interaction of Other Mutated Proteins with MAPKs

FGFR1, fibroblast growth factor receptor 1, interacts with MAPK3 with both its WT and mutant chains. It does not interact with the other MAPKs, it also does not have a structural preference for interacting with MAPK3. Protein of FGFR1 is a tyrosine kinase receptor [34].The involvement of this protein in VCI was not demonstrated previously but was mentioned in the context of other neurological disorders like schizophrenia and multiple sclerosis. This protein interacts with the MAPK/ERK pathway in a cancer system, regulating the epithelial-mesenchymal transition [35]. In our previous research we suggested an epithelial-mesenchymal transition in the blood brain barrier (BBB), caused by other two crosstalking pathways. As FGFR1 plays a protective role in BBB [36], we speculate it also can have a role in VCI regarding epithelial-mesenchymal transition.

The mutant chain of PARP1 (Poly ADP-Ribose Polymerase 1) has a conformational preference when interacting with MAPK14; it only interacts with one conformation. And also, another WT conformation of PARP1 preferentially binds to another WT MAPK14. PARP1’s has a preference when interacting with MAPK3, that is, it only interacts with one WT chain. PARP1 does not interact with MAPK8 with any of its conformations. Therefore, one WT chain of PARP1 has a structural preference for MAPK3, and the mutant chain prefers binding MAPK14. PARP1 is crucial for DNA repair and was related to neurodegenerative diseases previously[37].

AKT1(Protein Kinase B)’s mutant chain can interact with all the MAPKs, so it does not have a preference in the MAPKs. Two WT structures of AKT also do not have a structural preference for the MAPKs, as it interacts with all of them. The WT 1UNP chain A structure, on the other hand, has a structural preference for MAPK14 and MAPK8. This conformation only interacts with the one conformation of MAPK14, and one conformation of MAPK8. This points out that WT confirmation of AKT1 has structural preference when binding to MAPKs. AKT1 is a signaling molecule playing a role in the PI3K-Akt pathway. It was related to neurodegenerative diseases previously [38].

TP53’s mutant 3Q01 chain A interacts with all the given MAPKs. It interacts with one conformation of MAPK8; and three conformations of MAPK14. This means that this mutant conformation of TP53 has structural preference when interacting with MAPKs. Additionally, TP53’s mutant conformation 3Q05 chain B has a more selective interaction as it only interacts with MAPK14; and this interaction is present with two conformations of MAPK14. Regarding the WT interactions of TP53, chain A of 1OLG only interacts with MAPK3 and chain A of 2LGC of MAPK14. Showing a selectivity in terms of MAPKs and also in the conformation of MAPK14. The chain A of 5HP0 chain shows a selectivity towards MAPK14, such that it only interacts with one of its conformations. The D chain of 7BWN conformation of TP53 interacts with all MAPKs but has a structural preference for MAPK14 and MAPK8. Also, two other WT conformations of TP53, have a selective interaction with a conformation of MAPK8. These demonstrate that TP53 can interact with MAPKs through its many structural conformations, and it has unique preferences when interacting with MAPKs. We can therefore state that TP53 has a selective interaction with MAPKs with both its mutant and WT structures. TP53, p53 protein, takes place in many processes especially related to apoptosis, and cell cycle regulation. The interaction between MAPK3, MAPK8, and MAPK14 was previously discussed in the literature in the context of cancer. Here these kinases phosphorylate and activate the p53 protein [39]. Furthermore, it was suggested to have important functions in neurodegeneration [40]. Nevertheless, its interaction with MAPKs having an impact on VCI, was not directly suggested or demonstrated in literature to our knowledge. We suggest that the conformational preference when interacting with MAPKs can be a factor related to VCI.

## Methods

### 1. Selecting Disease Phenotypes related to Cardiovascular and Cognitive Diseases and Constructing Global Networks

Our previous work focused on finding novel interactions of VCI by building a Global Network (GN) [41]. For this, A disease-phenotype taxonomy was developed, and the CVD and CD phenotypes were chosen based on the number of seed proteins provided by GUILDify web server 2.0 [42]. We chose Atherosclerosis, Atrial Fibrillation, Hypertension, Myocardial Ischemia, and Stroke for CVD, and Dementia and Impaired Cognition for CD. Additionally, we created the network of OS because it is an important mechanism for the interaction between CVD and CD.

To understand the crosstalk we constructed two GNs, one for each tissue, one for the artery and one for brain tissue. These GNs are the result of merging the eight networks.

### 2. Weighted Average approach to find Hubs in the Global Networks

In order to find the hub proteins of the GNs, we performed a scoring analysis. For each protein in a GN we got the maximum CVD, CD score, the OS score and the betweenness centrality from their relevant GN. We ensured the scores are scaled. Later by initiating with equal weights we performed XGBoost. We compared the results to random weights. The final weight for each score was used to calculate a weighted score for each protein. We cut the scores from a threshold of score 1.

### 3. Including GWAS to hub proteins and finding their interactors

In order to study the interactions, we investigated the interactors of the hub proteins from the GNs, and included them along with the statistically significant genes coming from GWAS data. GWAS data was extracted using the API of EBI. The GWAS genes of the following traits were extracted. Atherosclerosis: EFO_0003914, Stroke: EFO_0000712, Atrial fibrillation: EFO_0000275, Hypertension: EFO_0000537, Myocardial Ischemia: EFO_1001375, Dementia: HP_0000726, Cognitive Impairment: HP_0100543, Vascular dementia: EFO_0004718. [43].

### 4. Creating A Protein-Protein Interaction Network with hubs and GWAS genes

To discover potentially new interactions affecting VCI, we created a PPI network with all the hubs, their first shell interactors, and GWAS genes using STRINGdb[44]. After downloading the network we used MCODE [45] with Cytoscape [46] to find the clusters of this network. MCODE settings are used as default (degree cutoff:2, node score cutoff:0.2, k-cut:2, maximum depth:100): The PPI network of the first cluster (or module) (highest MCODE score) was used for further analysis and its structural PPI Networks was built.

### 5. Constructing Structural Protein-Protein Interaction Network

Each protein in the PPI network’s PDB structure was identified and downloaded. Since PDBs of a protein can be redundant, it is necessary to group similar PDB structures together. Each cluster for a protein represents an alternative conformation of that protein and a representative PDB is chosen for each cluster. Sequence identity and structural similarity are used for clustering. Furthermore, the clustering process excludes proteins with fewer than 30 residues. Two PDB structures are considered to be in the same cluster if they share at least 95% of their amino acid sequences and have an RMSD value less than 2 A°. Finally, the structural PPI network (PDB-PDB network) was constructed with a set of scripts which was used as an input to protein docking with PRISM. The locations of the proteins were taken from UniProt [47]

### 6. Protein-Protein Interaction Modelling with PRISM and Mutation Analysis

PRISM is a tool for predicting interactions between proteins with template-based modeling of these interactions and protein structural analysis [3]. The creation of structural PPI takes place using PRISM.

The mutations of the relevant proteins were collected from UniProt [47] and ClinVar [48]. The effects of the mutations on the structural PPI were then examined by mapping the mutations to the PRISM modelings. In this research, only single nucleotide variations were taken into account.

We extracted the interaction localization information from the UniProt REST API. If two proteins are labeled in the same compartment, they are said to be colocalized.

## Acknowledgements

This project has been funded in part by TUBITAK Research Grant No: 220N252 and European Research Area Net (ERANET) ERA-CVD_JTC2020-015.

## Author Contributions

M.E.Z., S.S., A.G., and O.K. designed and conceptualized the project. M.E.Z. and S.S. analyzed data, prepared tables and figures. M.E.Z., S.S., A.G., O.K. wrote and edited the manuscript. All of the authors reviewed and approved the final manuscript.

## Data Availability

The datasets generated during and/or analyzed during the current study are available from the corresponding author on reasonable request.

## References

1. M, D. and L. D, Vascular Cognitive Impairment. Circulation research, 2017. 120(3).

2. EE, S., Clinical presentations and epidemiology of vascular dementia. Clinical science (London, England : 1979), 2017. 131(11).

3. A, B., et al., PRISM: a web server and repository for prediction of protein-protein interactions and modeling their 3D complexes. Nucleic acids research, 2014. 42(Web Server issue).

4. SM, R., et al., Nuclear receptor corepressor SMRT regulates mitochondrial oxidative metabolism and mediates aging-related metabolic deterioration. Cell metabolism, 2010. 12(6).

5. A, S., et al., c-Jun N-terminal kinase regulates soluble Aβ oligomers and cognitive impairment in AD mouse model. The Journal of biological chemistry, 2011. 286(51).

6. W, Z., et al., Loss of function of NCOR1 and NCOR2 impairs memory through a novel GABAergic hypothalamus-CA3 projection. Nature neuroscience, 2019. 22(2).

7. SK, L., et al., Silencing mediator of retinoic acid and thyroid hormone receptors, as a novel transcriptional corepressor molecule of activating protein-1, nuclear factor-kappaB, and serum response factor. The Journal of biological chemistry, 2000. 275(17).

8. Y, D., et al., Vascular CXCR4 Limits Atherosclerosis by Maintaining Arterial Integrity: Evidence From Mouse and Human Studies. Circulation, 2017. 136(4).

9. Q, H., et al., A review of the role of cav-1 in neuropathology and neural recovery after ischemic stroke. Journal of neuroinflammation, 2018. 15(1).

10. LW, B., et al., CXCR4 involvement in neurodegenerative diseases. Translational psychiatry, 2018. 8(1).

11. C, B., et al., Involvement of caveolin-1 in neurovascular unit remodeling after stroke: Effects on neovascularization and astrogliosis. Journal of cerebral blood flow and metabolism : official journal of the International Society of Cerebral Blood Flow and Metabolism, 2020. 40(1).

12. R, M., et al., Novel roles for the E3 ubiquitin ligase atrophin-interacting protein 4 and signal transduction adaptor molecule 1 in G protein-coupled receptor signaling. The Journal of biological chemistry, 2012. 287(12).

13. M, G.-H., R. R, and S. DF, Interactions of ErbB4 and Kap1 connect the growth factor and DNA damage response pathways. Molecular cancer research : MCR, 2010. 8(10).

14. X, C., et al., Mutant p53 in cancer: from molecular mechanism to therapeutic modulation. Cell death & disease, 2022. 13(11).

15. MI, E.-G., et al., A Review of HER4 (ErbB4) Kinase, Its Impact on Cancer, and Its Inhibitors. Molecules (Basel, Switzerland), 2021. 26(23).

16. AK, A., et al., Activation of p53 transcriptional activity by SMRT: a histone deacetylase 3-independent function of a transcriptional corepressor. Molecular and cellular biology, 2014. 34(7).

17. C, H., et al., Interactome Mapping Provides a Network of Neurodegenerative Disease Proteins and Uncovers Widespread Protein Aggregation in Affected Brains. Cell reports, 2020. 32(7).

18. S, G., et al., p53 amyloid formation leading to its loss of function: implications in cancer pathogenesis. Cell death and differentiation, 2017. 24(10).

19. F, C., et al., p53 is regulated by and regulates members of the gamma-secretase complex. Neuro-degenerative diseases, 2010. 7(1-3).

20. KM, F., et al., P53 aggregation, interactions with tau, and impaired DNA damage response in Alzheimer’s disease. Acta neuropathologica communications, 2020. 8(1).

21. RJ, O.B. and W. PC, Amyloid precursor protein processing and Alzheimer’s disease. Annual review of neuroscience, 2011. 34.

22. SM, G., et al., Amyloid angiopathy-related vascular cognitive impairment. Stroke, 2004. 35(11 Suppl 1).

23. A, T., et al., Amyloid burden, neuroinflammation, and links to cognitive decline after ischemic stroke. Stroke, 2014. 45(9).

24. SJ, N., et al., Blepharospasm in familial AD secondary to an APP mutation (V715M). Acta neurologica Belgica, 2014. 114(4).

25. K, A., et al., Unusual phenotypic alteration of beta amyloid precursor protein (betaAPP) maturation by a new Val-715 --> Met betaAPP-770 mutation responsible for probable early-onset Alzheimer’s disease. Proceedings of the National Academy of Sciences of the United States of America, 1999. 96(7).

26. HK, P., et al., Identification of PSEN1 and APP gene mutations in Korean patients with early-onset Alzheimer’s disease. Journal of Korean medical science, 2008. 23(2).

27. JB, K., et al., Novel Leu723Pro amyloid precursor protein mutation increases amyloid beta42(43) peptide levels and induces apoptosis. Annals of neurology, 2000. 47(2).

28. EV, B., et al., Familial L723P Mutation Can Shift the Distribution between the Alternative APP Transmembrane Domain Cleavage Cascades by Local Unfolding of the E-Cleavage Site Suggesting a Straightforward Mechanism of Alzheimer’s Disease Pathogenesis. ACS chemical biology, 2019. 14(7).

29. C, W., et al., Mitogen-activated protein kinase: conservation of a three-kinase module from yeast to human. Physiological reviews, 1999. 79(1).

30. B, C. and N. AR, Diversity and versatility of p38 kinase signalling in health and disease. Nature reviews. Molecular cell biology, 2021. 22(5).

31. W, Z. and L. HT, MAPK signal pathways in the regulation of cell proliferation in mammalian cells. Cell research, 2002. 12(1).

32. X, Z., et al., P38 activation mediates amyloid-beta cytotoxicity. Neurochemical research, 2005. 30(6-7).

33. SD, Y., et al., Cellular cofactors potentiating induction of stress and cytotoxicity by amyloid beta-peptide. Biochimica et biophysica acta, 2000. 1502(1).

34. S, D., et al., Fibroblast Growth Factor Receptors (FGFRs): Structures and Small Molecule Inhibitors. Cells, 2019. 8(6).

35. Y, H., et al., Regulation of brachyury by fibroblast growth factor receptor 1 in lung cancer. Oncotarget, 2016. 7(52).

36. J, C., et al., FGF21 Protects the Blood-Brain Barrier by Upregulating PPARγ via FGFR1/β-klotho after Traumatic Brain Injury. Journal of neurotrauma, 2018. 35(17).

37. ML, H., et al., Poly (ADP-ribose) polymerase 1 and neurodegenerative diseases: Past, present, and future. Ageing research reviews, 2023. 91.

38. A, G., et al., The PI3K-AKT pathway: A plausible therapeutic target in Parkinson’s disease. Experimental and molecular pathology, 2023. 129.

39. GS, W., The functional interactions between the p53 and MAPK signaling pathways. Cancer biology & therapy, 2004. 3(2).

40. Jr, C., et al., Role of p53 in neurodegenerative diseases. Neuro-degenerative diseases, 2012. 9(2).

41. Zeylan, M.E., et al., Revealing Shared Proteins and Pathways in Cardiovascular and Cognitive Diseases Using Protein Interaction Network Analysis. 2023.

42. J, A.-P., et al., GUILDify v2.0: A Tool to Identify Molecular Networks Underlying Human Diseases, Their Comorbidities and Their Druggable Targets. Journal of molecular biology, 2019. 431(13).

43. J, M., et al., Modeling sample variables with an Experimental Factor Ontology. Bioinformatics (Oxford, England), 2010. 26(8).

44. D, S., et al., The STRING database in 2021: customizable protein-protein networks, and functional characterization of user-uploaded gene/measurement sets. Nucleic acids research, 2021. 49(D1).

45. Bader, G.D. and C.W. Hogue, An automated method for finding molecular complexes in large protein interaction networks. BMC Bioinformatics, 2003. 4(1): p. 1–27.

46. P, S., et al., Cytoscape: a software environment for integrated models of biomolecular interaction networks. Genome research, 2003. 13(11).

47. Consortium, U., UniProt: the Universal Protein Knowledgebase in 2023. Nucleic acids research, 2023. 51(D1).

48. MJ, L., et al., ClinVar: improvements to accessing data. Nucleic acids research, 2020. 48(D1).

