## Supplementary Information for "Understanding Molecular Links of Vascular Cognitive Impairment: Selective Interaction between Mutant APP, TP53, and MAPKs"

FIGURES

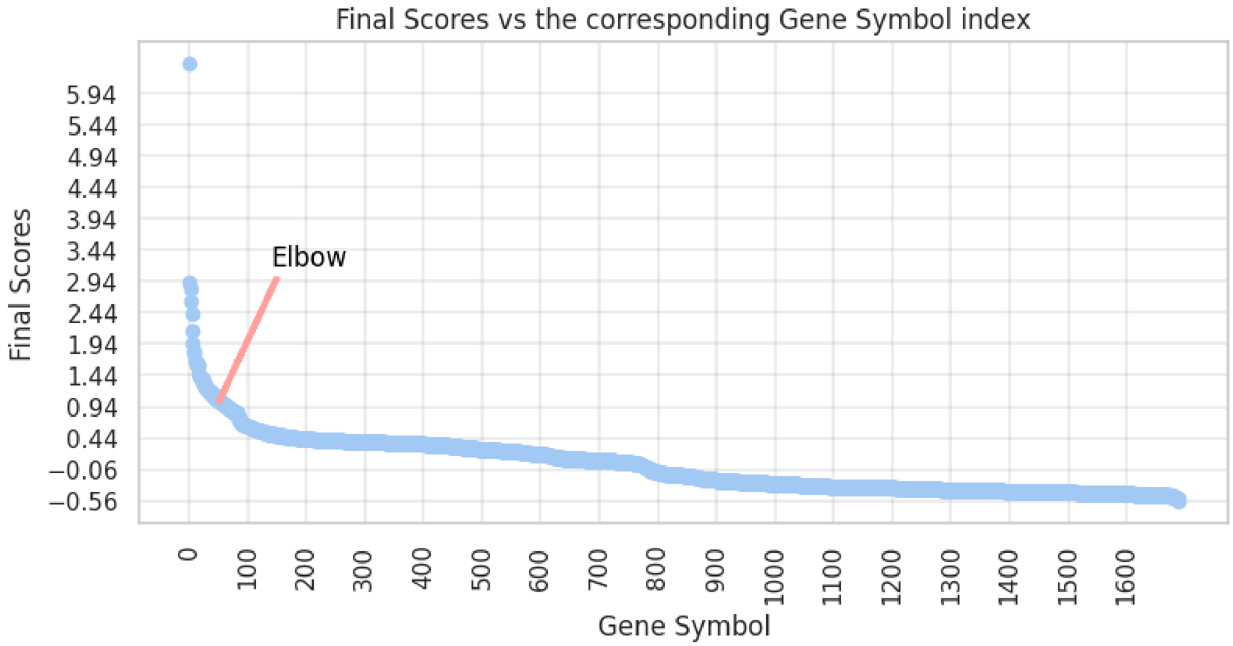


**Supplementary Figure 1:** Score distribution for the genes in the brain tissue related global network. Elbow at score 1 in which there are 57 proteins with a higher score than 1.

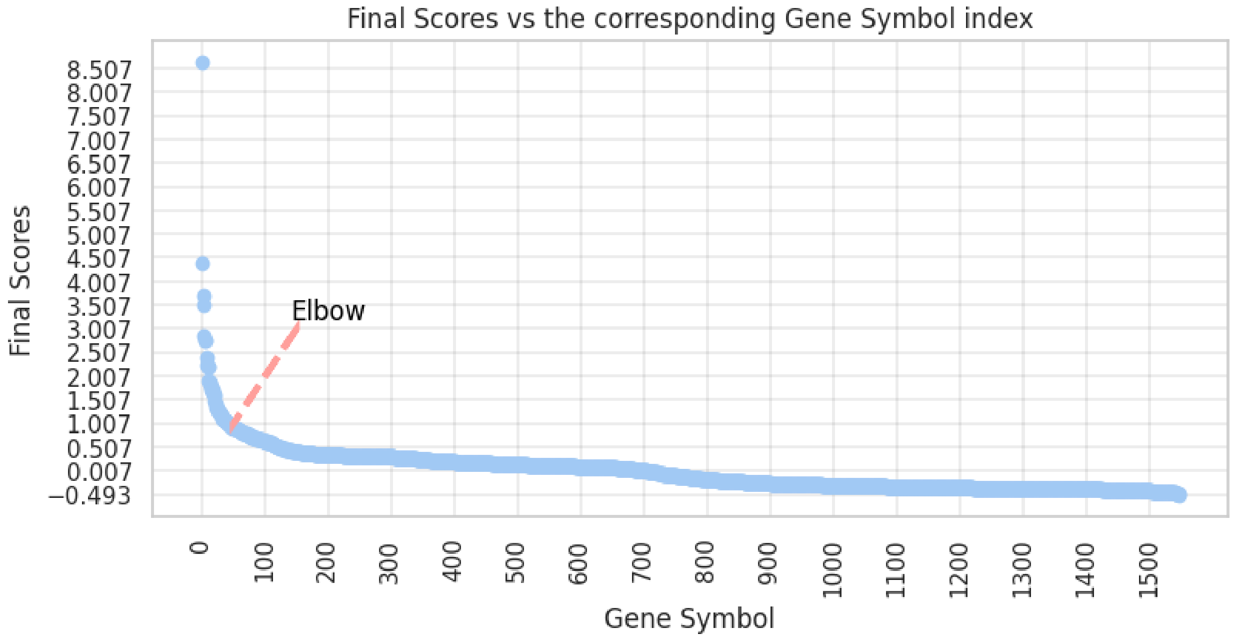


**Supplementary Figure 2:** Score distribution for the genes in the artery tissue related global network. Elbow at score 1 in which there are 41 proteins with a higher score than 1.
